# The genealogical decomposition of a matrix population model with applications to the aggregation of stages

**DOI:** 10.1101/067793

**Authors:** François Bienvenu, Erol Akçay, Stéphane Legendre, David M. McCandlish

## Abstract

Matrix projection models are a central tool in many areas of population biology. In most applications, one starts from the projection matrix to quantify the asymptotic growth rate of the population (the dominant eigenvalue), the stable stage distribution, and the reproductive values (the dominant right and left eigenvectors, respectively). Any primitive projection matrix also has an associated ergodic Markov chain that contains information about the genealogy of the population. In this paper, we show that these facts can be used to specify any matrix population model as a triple consisting of the ergodic Markov matrix, the dominant eigenvalue and one of the corresponding eigenvectors. This decomposition of the projection matrix separates properties associated with lineages from those associated with individuals. It also clarifies the relationships between many quantities commonly used to describe such models, including the relationship between eigenvalue sensitivities and elasticities. We illustrate the utility of such a decomposition by introducing a new method for aggregating classes in a matrix population model to produce a simpler model with a smaller number of classes. Unlike the standard method, our method has the advantage of preserving reproductive values and elasticities. It also has conceptually satisfying properties such as commuting with changes of units.

## 1. Introduction

Many simple models in population biology take the following form: a non-negative vector gives the current abundances of types within the population; then, to determine the abundances of types at some future time, one multiplies this vector by a non-negative matrix capturing the interconversion and reproductive rates of the types. Examples include models of deterministic mutation-selection balance in population genetics (where the types correspond to genotypes, Nagylaki, 1992 Chapter 2; Bürger, 2000 Chapter 3) and models of spatially structured populations (where the types correspond to demes, Rousset, 2004). The most common use of such models is in the ecological and demographic literature, where the types correspond to age ranges or developmental stages. In this last context, such models are commonly known as “matrix population models” and they play a critical role in both ecological theory and applications to population management (Caswell, 2001).

In the ecological or demographic context, the entries in the update or projection matrix are typically estimated based on observations from some natural population (Salguero-Gómez et al., 2015; Salguero-Gómez et al., 2016). To better understand the dynamics of the population, one then calculates various descriptors of the resulting model such as the asymptotic growth rate of the population, the generation time, the asymptotic distribution of type frequencies, etc. (for a more complete list, see e.g. Cochran and Ellner, 1992; Caswell, 2001). Here, we provide a method to move in the opposite direction: given certain descriptors of the population, we construct the corresponding projection matrix. Besides providing a means to construct projection matrices with specified properties, our method provides a unifying perspective on the theory of matrix population models by clarifying the relationships between various commonly used descriptors.

The key idea is that any matrix population model is completely determined by the specification of (1) its asymptotic growth rate, (2) its stable stage distribution and (3) a Markov chain describing the sequence of classes visited when we consider the lineages of individuals within the population. While this viewpoint is perhaps implicit in the classical literature (Demetrius, 1974, 1975; Tuljapurkar, 1982, 1993), its power has not been sufficiently appreciated because the strength of the connections between this genealogical Markov chain and other population descriptors has only recently come to light. In particular, recent work has revealed that certain hitting times on this genealogical Markov chain determine the generation time (Bienvenu and Legendre, 2015; Lehmann, 2014a), while the asymptotic frequencies of the transitions of this Markov chain give the elasticities of the asymptotic growth rate with respect to the entries of the projection matrix (Bienvenu and Legendre, 2015). Since an ergodic Markov chain is uniquely specified by the asymptotic frequencies of its transitions, this means that if we specify the asymptotic growth rate, stable stage distribution and matrix of eigenvalue elasticities, we can immediately write down the unique projection matrix with these desired characteristics.

This construction provides a great deal of clarity, particularly concerning the interpretation and biological meaning of eigenvalue elasticites. Indeed, merely recognizing that the matrix of elasticities is given by the asymptotic transition frequencies of the genealogical Markov chain makes several facts obvious that are otherwise rather mysterious from a classical perspective (Bienvenu and Legendre, 2015). For instance, one can show that the total of the entries of the elasticity matrix must sum to one by either direct calculation (de Kroon et al., 1986) or by an appeal to Euler’s Theorem for homogeneous functions (Mesterton-Gibbons, 1993). However, recognizing the elasticities as the asymptotic transition frequencies of a Markov chain make it obvious that they sum to one, since the asymptotic frequencies of the transitions form a probability distribution (the chain must always transition from one state to another). Similarly, the row sums of the matrix of elasticities equal its column sums (van Groenendael et al., 1994) due to the simple fact that at stationarity the probability of arriving in a state must equal the probability of exiting that state. Furthermore these row and column sums are just the class reproductive values, which when appropriately normalized are themselves just the asymptotic frequencies of the classes visited by the genealogical Markov chain.

The present work shows how a Markov chain perspective can be carried further to illuminate other aspects of the theory of matrix population models. For instance, it is helpful to classify descriptors of the matrix population models in terms of their dependencies on the triple of growth rate, stable stage distribution, and genealogical Markov chain: in our parametrization, elasticities depend only on the genealogical Markov chain, whereas the sensitivities of the asymptotic growth rate to perturbations in the entries of the projection matrix do not depend on the asymptotic growth rate but do depend on both the genealogical Markov chain and the stable stage distribution. Similarly, whereas matrix population models most frequently track the number of individuals in a given class, they can also be written in terms of other units such as the biomass present in each class. It turns out that specifying the stable stage distribution is equivalent to making a choice of units, so that, for example, the matrix of sensitivities depends on the choice of units whereas the matrix of elasticities does not. Indeed, the genealogical Markov chains arise by expressing the matrix population model in units of reproductive value, so that the choice of stable stage distribution can be fruitfully viewed as determining the conversion factor between reproductive value and number of individuals. That is, two models can have the same genealogical Markov chain and asymptotic growth rate but different stable stage distributions because of different choices concerning how reproductive value is packaged into individuals.

To demonstrate the power of this approach, we present a new solution to the problem of how to aggregate states in a matrix population model. This problem is important for two reasons. First, it has long been known that estimates of various population descriptors depend on the number of organismal states used in the matrix population model (Silvertown et al., 1993; Enright et al., 1995; Benton and Grant, 1999; Ramula and Lehtilä, 2005; Salguero-Gómez and Plotkin, 2010; Picard and Liang, 2014). As a result, when comparing matrix projection models of different species, the dimensionality of the projection matrix is sometimes reduced by aggregating or “collapsing” multiple states into one so that the dimensionality is the same for all species being compared (Enright et al., 1995; Salguero-Gómez and Plotkin, 2010). Second, because one needs to observe multiple transitions between pairs of classes to accurately estimate vital rates, there is a trade-off between error in estimating the vital rates and the degree of within-state heterogeneity that is neglected by the model (Vandermeer, 1978; Moloney, 1986; Caswell, 2001). Thus, some degree of collapsing necessarily arises in the construction of matrix population models, a defect which in part motivated the proposal of integral projection models (Easterling et al., 2000).

The standard method for collapsing states in matrix population models was proposed by Enright et al. (1995) and generalized by Salguero-Gómez and Plotkin (2010). It essentially assumes that the population is at its stable stage distribution and then aggregates a group of classes by considering what we would observe if we did not distinguish between classes within this collapsed group. Remarkably, this procedure preserves both the asymptotic growth rate and the stable stage distribution (Hooley, 2000; Salguero-Gómez and Plotkin, 2010). However, its effects on reproductive values and elasticities are poorly characterized and can be substantial (Enright et al., 1995; Benton and Grant, 1999; Ramula and Lehtilä, 2005; Salguero-Gomez and Plotkin, 2010; Picard and Liang, 2014).

Here we show that this behavior arises because the standard method, while preserving the stable stage distribution and asymptotic growth rate, fails to preserve the genealogical Markov chain. By applying our decomposition to the projection matrix, we propose a method wherein the stable stage distribution and genealogical Markov chain are collapsed separately and subsequently recombined to construct the collapsed projection matrix. This method optimally preserves reproductive values, the genealogical Markov chain, the matrix of elasticities, and the generation time in addition to the stable stage distribution and the asymptotic growth rate. The method is also independent of the units used to describe the population in the sense that, unlike the standard collapsing method, it commutes with changes of units. We return to the practical applicability of this new collapsing method in the Discussion.

## 2. Genealogical Markov chains associated with matrix population models

A matrix population model is given by a non-negative matrix **A** = (*a_ij_*). The model assumes that if there are *n_j_*(t) individuals in the population of class *j* at time *t*, these individuals will make a contribution of *a_ij_ n_j_*(*t*) individuals to the total number of individuals of class *i* at time *t* +1. That is, the dynamics of the population are governed by the matrix equation

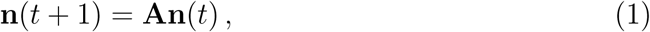
where *n*(*t*) = (*n_i_*(*t*)) is the vector giving the number of individuals in each class at time *t*.

While equation (1) describes the dynamics of the size and composition of the population, it is also sometimes useful to consider the sequence of classes occupied by a particular individual, its ancestors, and descendants. We begin by reviewing the features of two Markov chains that capture the dynamics along such lineages. These ideas are due to Demetrius (1974, 1975), and have been further exploited in Tuljapurkar (1982, 1993).

### 2.1 The backward chain, P

Suppose we want to know the probability that an individual observed in class i at time *t* comes from class *j*, which can have one of two meanings: if the individual was alive at time (*t* − 1), we want to know the probability that it was in class *j*; conversely, if the individual is a newborn at time *t*, we ask what is the probability that its mother was in class *j*. Let **A** = (*a_ij_*) be the population projection matrix and **n**(*t*) = (*n_i_*(*t*)) be the population vector. Since there are *n_i_*(*t*) individuals in class *i* at time *t, a_ij_* n*_j_*(*t* − 1) of which come from class *j*, this probability is given by

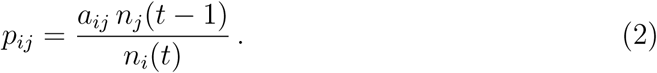
Now, let us assume that **A** is primitive, i.e. that there exists a non-negative integer *t* such that all the entries of **A***^t^* are strictly positive. In that case, the population structure will converge to a unique stable distribution given by the vector w whose entries sum to one and satisfying

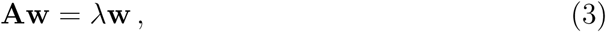
where λ, the greatest eigenvalue of **A**, gives the asymptotic growth rate of the population (Caswell, 2001). If we then assume that the population is at its stable stage distribution, **n**(*t*) ∝ **w**, we have *n_j_*(*t*−1)/*n_i_*(*t*) = *w_j_/*(λ*w_i_*) and Equation (2) becomes

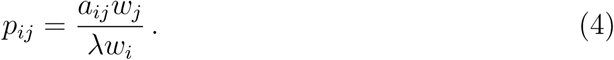
Because each individual in the population at time *t* is descended from some individual at *t* − 1, the rows of the matrix **P** = (*p_ij_*) sum to one, so that it is a Markov matrix. Furthermore, the associated Markov chain is ergodic due to the primitivity of **A**, since **P** and **A** have the same pattern of non-zero entries. This Markov chain allows one to track the classes visited by a lineage backwards in time, i.e., it records the (infinite) sequence of stages occupied by an individual and its ancestors.

The stationary probability distribution of **P**, which corresponds to the asymptotic proportion of time that this backwards-time lineage spends in each class, is given by **π**, the left eigenvector of P satisfying **π** = **πΡ**, whose entries have been scaled to sum to 1. Moreover,

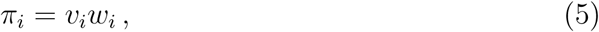
where **v**, the (row) vector of reproductive values satisfying **vA** = **λv**, has been scaled so that **vw** = 1. This product *v_i_w_i_* = *π_i_* is also known as the class reproductive value. Whereas the individual reproductive value *v_i_* gives the asymptotic contribution of an individual of class i to the total size of the population far in the future, the class reproductive value gives the asymptotic proportional decrease in the size of the population far in the future if we were to kill all class *i* individuals at the present time in a population at its stationary stage distribution. Thus, equation (5) shows that the asymptotic proportion of time a lineage spends in class *i* going backward in time, *π_i_*, is equal to the proportional reduction in long-term population-size due to a single culling event of all individuals in class *i* in an otherwise stationary population.

The matrix **P** also appears in several other contexts. In population-genetics, it is known as the “backward migration matrix” and plays a central role in the theory of evolution in class-structured populations where it is used to update the class-specific allele frequencies from one generation to the next (Bodmer and Cavalli-Sforza 1968, Taylor 1990, Rousset and Ronce 2004, Rousset 2004, pp. 190-192). In that literature, the key observation is that the allele frequency in class *i* in the current generation, *x_i_*(*t*), is the average of the allele frequencies for the other classes *j* in the previous generation weighted by the probability that an individual currently in class *i* was descended from an individual in class *j*, so that 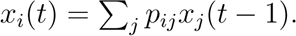.

In ecology, the importance of the Markov chain defined by **P** arises due to the close connection between the backwards-time Markov chain defined by **P** and the eigenvalue elasticities of the transition matrix. In particular, the frequencies of transitions between states in the stationary chain defined by **P** (i.e. transitions along the arcs of the life-cycle graph) are equal to the elasticities of the asymptotic growth rate λ to the entries of the projection matrix (Bienvenu and Legendre, 2015):

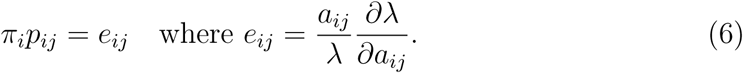
Many formal features of the matrix of elasticities **E** = (*e_ij_*), such as the fact that the entries sum to one (de Kroon et al., 1986) and the fact that the row sums equal the column sums (that is, 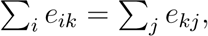, van Groenendael et al. 1994) are obvious once one recognizes the matrix of elasticities as the asymptotic transition frequencies of this genealogical Markov chain. At the same time the stationary distribution *π* (class reproductive values) gives the diagonal entries of the sensitivity matrix, i.e.

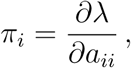
and the generation time is simply related to certain sums of the stationary transition probabilities *π_i_p_i_j* (Bienvenu and Legendre, 2015).

### 2.2 The forward chain, Q

The transition matrix **P** allows us to study how a lineage moves through the set of classes as we look backwards in time. We now define a corresponding process that describes the classes occupied by the descendants of a given individual as we look along its lineage forward in time. The following construction is based on Tuljapurkar (1982, 1993), with a straightforward generalization from Leslie matrices to arbitrary primitive projection matrices.

While looking at lineages going backwards in time was relatively simple because each individual has exactly one ancestor at any prior time, looking at lineages going forwards in time is more complex due to the branching nature of genealogies. Thus, to follow a lineage forwards in time requires decisions about which branches to follow. Most importantly, we need to ensure that we will not get trapped in a ‘dead-end’ by following a lineage that does not leave any descendants after a certain time.

We can do so by picking an individual in the distant future and identifying its ancestor at the present time. We then start from this ancestor and choose to follow the branches that lead to the individual that we picked in the distant future. Because far in the future the population is at its stationary stage distribution, when we trace this lineage backwards in time the sequence of classes visited is described by the Markov chain **P** of the previous section. Therefore, to describe it in forward time all we have to do is to consider the time-reversed Markov chain of **P**, i.e. the Markov chain **Q** whose probability transitions are given by

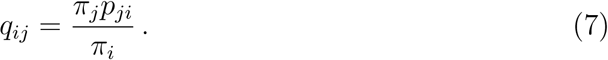
Equation (7) simply expresses the fact that, at stationarity, the probability of going from *i* to *j* in forward time is the same as the probability of going from j to i in backward time. Note that the ergodicity of **P** is required to ensure that *π_i_* > 0 in the definition of **Q**, and that **Q** is ergodic (since it has the same pattern of non-zero entries as **P**^T^). Finally, it is easy to check that 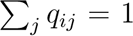 and that the stationary probability distribution of **Q** is **π**. For more on time-reversed Markov chains, see e.g. section 5.3 of Kemeny and Snell (1976).

Substituting for *p_ij_* and *π_i_* in equation (7), we get

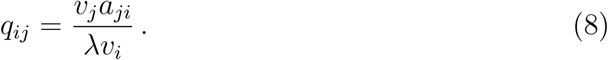
This expression has a straightforward biological interpretation, which can in fact be used to define **Q** without referring to **P**: first, recall that reproductive values give the contributions of individuals to the long-term growth of the population. Indeed,

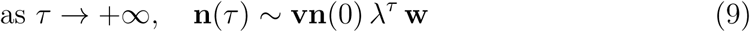
so each of the *n_i_*(0) individuals in class *i* at time 0 contributes *v_i_*λ^τ^ to the 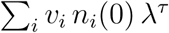 individuals in the population at time τ (a proof of equation (9) can be found in Chapter 1 of Seneta, 2006). Therefore, we can write down the probability that an individual at time τ ≫ *t* had its ancestor in class *j* at time *t* +1 given that its ancestor was in class *i* at time *t*: indeed, the ancestor at time *t* will leave *a_ji_* descendants in class *j* at time *t* + 1, each of them contributing *v_j_*λ^τ-(^*^t^*^+1)^ to the population at time τ. Thus, out of the *v_j_*λ^τ-1^ individuals left by the ancestor in class *i* at time *t, v_j_*λ^τ-(^*^t^*^+1)^*a_ji_* will be descended from an individual in class *j* at time *t* + 1. As a result,

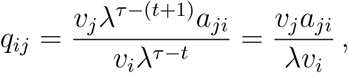
and we recover equation (8).

## 3. The genealogical decomposition

So far we have worked from a given projection matrix **A**, derived the corresponding Markov chains defined by **P** and **Q** and discussed some of the properties of these chains. This is in line with the most common applications of matrix population models where one estimates **A** from data and then computes various descriptors of the population using the entries of **A**. However, sometimes it might be useful to work in the opposite direction, for instance if we have observed the values of some descriptors and we want to know the set of all projection matrices that are compatible with the observed values (e.g. if we know the generation time and stable stage distribution, what values are possible for the unobserved vital rates?). Moreover, from a theoretical perspective it is useful to understand whether the value of one descriptor constrains the possible values of another, or alternatively if the two descriptors can be “chosen” arbitrarily, in the sense that for any pair of values we can construct a corresponding projection matrix.

In particular, given a forward or backward time genealogical Markov chain defined by a primitive matrix **P** or **Q**, we would like to construct a corresponding projection matrix **A** that exhibits the appropriate genealogy. To do this, first notice that **P**, **Q** and the matrix of elasticities E all uniquely determine each other, and each of these also determines the vector of class reproductive values **π**. In particular, **π** is the dominant left eigenvector of **P** and **Q** and is also given by the row and column sums of **E**, that is,

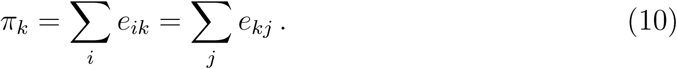
Given π, one can then convert between the entries of **P**, **Q**, and **E** using equations (6) and (8).

Once **P**, **Q** or **E** is specified, we can then construct a corresponding matrix **A** by solving for *a*_*ij*_ in equation (4):

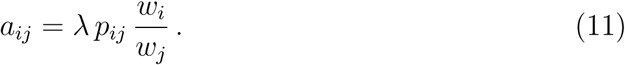
Thus, for a given choice of **P**, **Q** or **E** we can explicitly construct all possible compatible projection matrices by making a choice of stationary stage distribution w and asymptotic growth rate λ. Alternatively, we can choose the vector of reproductive values v and the growth rate λ since, once **P**, **Q** or **E** is given, w and v determine each other by the relations π*_i_* = *v_i_w_i_*. Assuming that the specified **w** (or **v**) and λ are strictly positive, the resulting **A** is primitive because it has the same pattern of non-zero entries as **P**, which is primitive. Note that these positivity conditions on **w** (or **v**) and λ do not represent a substantial constraint on our choice of **w** and λ because, by the Perron-Frobenius Theorem, these conditions hold for any primitive matrix **A**.

The previous paragraphs shows that it is possible to specify the **P** matrix independently of **w** and λ. But from an intuitive perspective, the fact that the genealogical dynamics can be decoupled from the relative abundances of the classes at stationarity is perhaps surprising. To see why this is the case, it is helpful to consider what happens to a matrix population model when we change the units that the model is expressed in. For instance, instead of tracking the number of individuals in each class we could track the amount of biomass in each class, where individuals within a class have uniform masses. Indeed, this might be quite useful for management purposes in the case of forests or fisheries where we might primarily be interested in yields rather than the number of individual organisms. Using units different from individuals might also be relevant in situations where it is hard to count individuals (e.g., plants forming thick mats where it is hard to isolate individuals but easy to count the number of fruits or to measure the area of the mat; colonies of microorganisms, etc) or in which we are interested in the number of gene copies and individuals in different classes have different ploidy. More generally, if *c_i_* is the number of new units in class *i* for each old unit in class *i* (e.g. *c_i_* is the mass in grams per individual of class *i*), then the entries of the matrix population model in the new units are given by

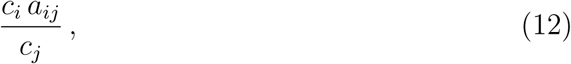
and the new stable stage distribution has entries proportional to *c_i_w_i_* and reproductive values (per unit) of *v_i_/c_i_*. Importantly, when we calculate the backward Markov chain we see that it is invariant to changes of units since

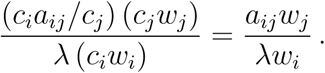
This corresponds to the simple fact that when tracing genealogies, it doesn’t matter whether we track where a given gram came from in the previous time step or where the individual containing that gram came from. The genealogical Markov chains are likewise independent of the asymptotic growth rate λ because changes in λ only change the absolute size of the population and not the proportions in each class. Because the stable stage distribution depends on the choice of units (picking a random gram is different than picking a random organism), by choosing the units appropriately, we can produce an arbitrary stable stage distribution while maintaining a fixed genealogical Markov chain.

The concept of changes of units also provides an additional perspective on the matrices **P** and **Q** in that these matrices arise when we express **A** in terms of its natural units. In particular, if we pick *c_i_* = *v_i_*, so that the dynamics of the population are expressed in units of reproductive value, then the new projection matrix is λ **Q**^T^ (see Equation 8; the transpose arises because we are following the convention that projection matrices act on column vectors and transition matrices act on row vectors) and the stable stage distribution is **π**, the vector of class reproductive values. Thus, another way of viewing the stable stage distribution of the original projection matrix **A** is as a way of partitioning a given amount of class reproductive value, π*_i_*, to a certain number of individuals. For a fixed value of π*_i_* = *w_i_v_i_*, one can choose to have either a large number of individuals each with a small reproductive value (large *w_i_*, small *v_i_*) or a small number of individuals each with a relatively large reproductive value (large *v_i_*, small *w_i_*). This flexibility is another way of understanding why a fixed set of genealogical dynamics is compatible with an arbitrary observed stable distribution of the population across classes.

### 3.1 Descriptors and their relations

The above discussion helps clarify the relationships between several commonly used descriptors of matrix population models. In particular, it is helpful in understanding the relationship between the sensitivities and elasticities of λ with respect to the *a_ij_*. While the elasticities are completely determined by the genealogical Markov chain, the sensitivities

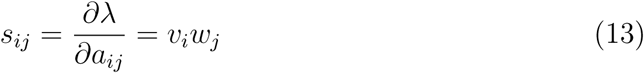
depend on both the genealogical Markov chain and the choice of units. This makes sense because elasticities measure the effects of multiplicative perturbations which are independent of the choice of units (doubling a rate results in twice as much output, whether the input and output are measured in individuals or grams), whereas sensitivities measure the effects of additive perturbations, which depend on the choice of units (producing an extra fraction of an individual per individual is physically different than producing an extra fraction of a gram per gram if individuals of different classes do not have the same mass). The exception to this dependence on the choice of units are the diagonal entries of the matrix of sensitivities *s_ii_* = *v_i_w_i_* = π*_i_*, since the aii are invariant to the choice of units and the sensitivities are equal to the class reproductive values. Interestingly, the sensitivities only depend on the genealogical Markov chain through its stationary distribution (i.e. through the class reproductive values n_i_). Thus, sensitivities contain information not contained in the elasticities in that they reflect a choice of units, but elasticities contain information not contained in the sensitivities in that they reflect the specific paths that lineages tend to take through the life cycle and not just the fraction of time lineages take in each stage.

Another important set of descriptors are the full set of eigenvalues of the projection matrix. These are determined by λ together with the genealogical Markov chain and are independent of the choice of units, reflecting the more general fact that the eigenvalues of a matrix are independent of the basis it is expressed in. Importantly, the genealogical Markov chain alone determines the ratios between eigenvalues (e.g. the damping ratio), which explains why knowledge of the genealogical Markov chain alone is sufficient to understand the approach to stationarity (Tuljapurkar, 1982, 1993).

### 3.2 Two parametrizations of 2 × 2 models

To illustrate the discussion above, we work out the (**P**, **w**, λ) parametrization in the case of 2 × 2 models. Any 2 × 2 matrix model can be parametrized by 4 non-negative numbers *a, b, c* and *d* through the projection matrix

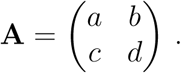
Though these four parameters usually have straightforward individual-based interpretations, in the general case the link between them and the descriptors of the population is not immediate.

To write out the (**P**, **w**, λ) parametrization, we start by noting that any ergodic Markov matrix P can be parametrized by two numbers *p* and *q* in ]0,1], where either *p* or *q* can be equal to 1, but not both:

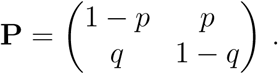
Similarly, any stable distribution vector w is parametrized by a single number *x* > 0:

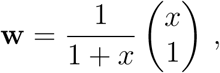
Choosing λ as the fourth parameter and using equation (11), we get the following parametrization for **A**:

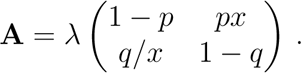
Although it looks more complicated than the (*a, b, c, d*) parametrization, this parametrization has the advantage of taking important population-level descriptors such as the growth rate λ or the ratio *x* = *w_1_/w_2_* directly as parameters. Similarly, the elasticity and sensitivity matrices of these models are given by

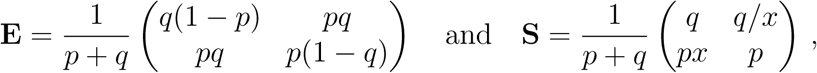
and the eigenvalues are λ and λ(1 − *p* − *q*) so that the damping ratio is 1/|1 − *p* − *q|* (which, as noted previously, depends only on *p* and *q*).

In principle, the (**P**, **w**, λ) parametrization could provide an alternative way to build a projection matrix from field data (for instance, in the 2 x 2 case, λ can be estimated by *n*(*t* + 1)/*n*(*t*); *x* by *n*_1_(*t*)*/n*(*t*)*; p* by the fraction of newborns and 1 − *q* by the fraction of individuals in class 2 that were already in class 2 in the previous year). However, our results are likely to be most useful in theoretical studies of matrix populations models: for instance, by providing a way to specify “random” projection matrices with prescribed descriptors (e.g., to test a conjecture), or to compare models (e.g., see what the fertilities and survival probabilities must be to have a model with the same stationary distribution and elasticities but a higher growth rate). As an illustration of the utility of our approach, we turn now to our main application of the genealogical decomposition: reducing the number of states in a matrix population model while optimally preserving the properties of the original model.

## 4. Application: collapsing states

We have suggested that in some cases it might be more enlightening to think of a matrix population model as a (genealogy, partitioning of class reproductive value, growth rate) ≡ (P, w, λ) triple rather than as a population projection matrix. We illustrate this by proposing a new method for aggregating a set of states in a matrix population model into a single state. As discussed in the Introduction, this is an important practical problem when trying to compare projection matrices of different sizes, because the dimensionality of the matrix population model is known to affect the estimates of various descriptors (Silvertown et al., 1993; Enright et al., 1995; Benton and Grant, 1999; Ramula and Lehtilä, 2005; Salguero-Gómez and Plotkin, 2010; Picard and Liang, 2014).

### 4.1 Individualistic collapsing

The most natural way to collapse a model is to put oneself in the shoes of the experimenter building the model, and then simply disregard the now superfluous distinctions between individuals in the collapsed classes. To do so, we use the model that we want to collapse to “simulate” the population dynamics. Then, we build the collapsed model by counting individuals in terms of the collapsed classes and interpreting the entries of the matrix as per capita contributions - that is, *a_kl_* is the number of individuals of class *k* that come from class *l* divided by the number of individuals that were in class *l*. As a result, the total contribution of collapsed class *j* to collapsed class *i* is obtained by counting the number of individuals coming from any class *l* collapsed to *j* that are present in any class *k* collapsed to *i*. Dividing by the total number of individuals in collapsed class *j*, we get the per-capita contribution of collapsed class *j* to collapsed class *i*:

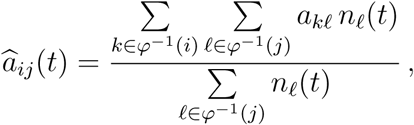
where the *a_kl_,*S are the entries of the original model and the 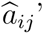s those of the collapsed model, and *φ* is the collapsing function, which is defined by *φ*(*k*) = *i* if and only if class *k* corresponds to class *i* in the new model. *φ*^−1^(*i*) denotes the preimage of *i* by *φ*, i.e. the set of classes in **A** that are collapsed to *i* in 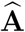. For simplicity, from now on we will note *k* ⊂ *i* for *k* ∈ *φ*^-1^(*i*), the justification for this notation being that *k* can be thought of as a subclass of *i*.

If we want this collapsing to be independent of the composition of the population, we can assume that the population is at its stable stage distribution, w, in which case we have

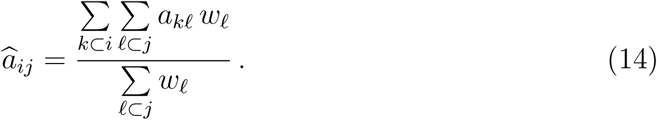
We call this method *individualistic collapsing* and hereafter refer to 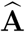 as the individualistic model and to **A** as the original (or extended) model. We use hats to denote any quantity associated with the individualistic model.

Individualistic collapsing is the standard method used for collapsing projection matrices. It was introduced by Enright et al. (1995) and later generalized by Salguero-Gómez and Plotkin (2010). Its properties, which we now review, have been studied by Hooley (2000).

We start by noting that individualistic collapsing preserves the asymptotic growth rate and the stable stage distribution. Indeed, let

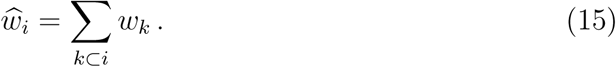
Then,

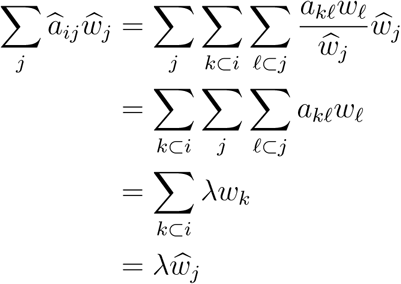
Moreover, if **A** is primitive, then clearly so is 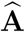 and thus the Perron-Frobenius theorem ensures that there is only one (that is, up to a multiplicative constant) right eigenvector of 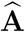 with only positive entries (Seneta, 2006). Since this is the case of the vector w defined in (15), it is the dominant right-eigenvector of 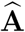 and λ is the associated eigenvalue.

What about reproductive values? The classic interpretation of these quantities suggests that we should have

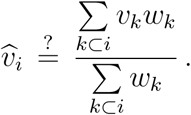
However, unless 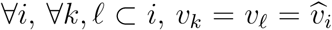, that is, if the classes collapsed together have the same reproductive value (in which case we clearly have 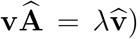, this candidate is not a left-eigenvector of 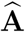. Furthermore, reproductive values behave in a highly unintuitive way under individualistic collapsing, as illustrated by the fact that collapsing a set of classes can change the reproductive values of other classes that are not collapsed.

Another shortcoming of individualistic collapsing is that it does not commute with the construction of the genealogical matrices, in the sense that the descriptors of the Markov chain **P** constructed from **A** are not compatible with those of the Markov chain 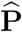 constructed from 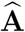. Take for instance the stationary probability distributions, π and 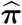: clearly, since the time spent in class *i* by a lineage is the sum of the time spent in each of the subclasses of i, 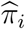 should be equal to 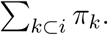 However, this is not the case, since, unless 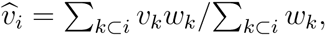

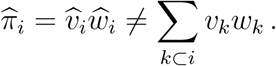
As a result of this, the biological descriptors which depend on **P**, such as the generation time, will not be the same for **A** and 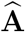. Most importantly, the elasticities are affected: since they quantify the relative change in λ in response to a multiplicative perturbation of the entries of the matrix, we should expect the elasticity of the collapsed entry 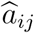 to be the relative change in λ when all the corresponding entries in **A** are subjected to the same multiplicative perturbation. Thus, if we write

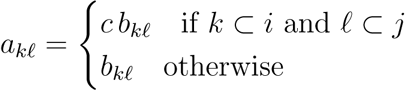
and evaluate around *c* = 1 (which implies 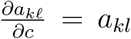 if *k* ⊂ *i* and ℓ ⊂ *j*, and 0 otherwise), then we should have

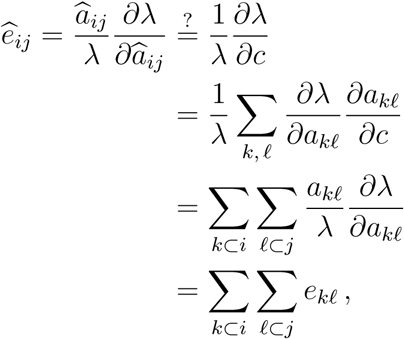
i.e. the elasticity of λ to 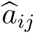 should be the sum of the elasticities of *λ* to the corresponding entries of **A**, in accordance with the interpretation of the elasticities as asymptotic frequencies of traversal of the arcs. However, this is not the case with individualistic collapsing, since 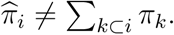.

Finally, note that individualistic collapsing does not commute with changes of units, in the sense that if we define the collapsed conversion factors naturally as the weighted average within each collapsed class, i.e.

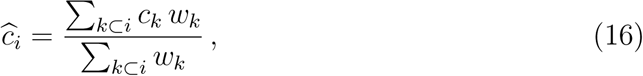
then we see that switching to a given unit, collapsing and then switching back to the original unit gives a model different from the one obtained by collapsing alone. In other words, individualistic collapsing depends on the units the model is expressed in.

### 4.2 Genealogical collapsing

Given the limitations of individualistic collapsing, we look for an alternative method. Our framework suggests collapsing the (genealogy, partitioning of reproductive value, growth rate) triple rather than the projection matrix **A**, as illustrated by figure 1. The idea behind this approach is that we will be able to choose how we collapse each of the separate components of the triple in order to preserve specific properties. This construction can be done using either of **P**, **Q** or **E** for the genealogical part of the triple, and either of w or v for the distribution of class reproductive value among the classes of the model - all these possibilities are equivalent and will give the same result. Here we use (**P**, **w**, λ).

**Figure 1:**
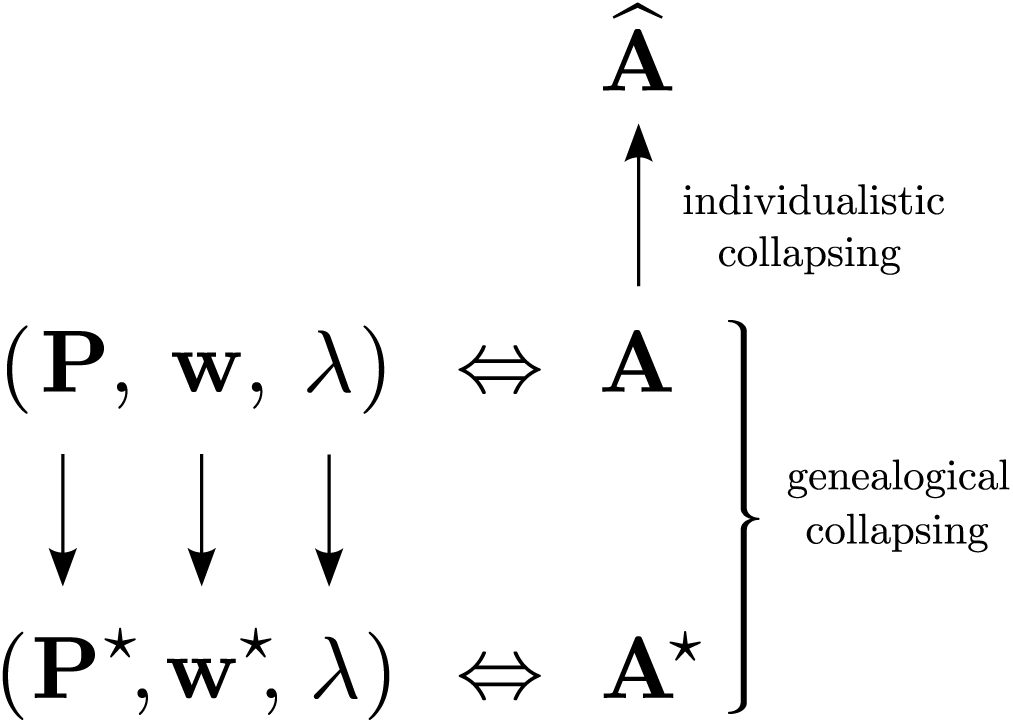
Graphical comparison of the two collapsing strategies.

If we want to preserve the asymptotic growth rate and the stable stage distribution, then w and λ should be collapsed according to

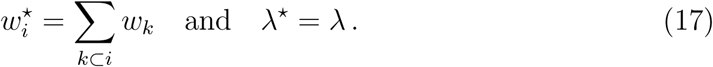
Therefore, all we have to do is specify how to collapse **P**. Some precaution is needed here as, in the general case, it is impossible to collapse a Markov chain – in the sense that the process we obtain by aggregating the states of a Markov chain is not a Markov chain. To see this, consider a Markov chain that goes from state i to state *i* + 1 and then from *i* + 1 to *i* + 2 with probability 1. If states *i* and *i* + 1 are to be collapsed together, the resulting system should spend exactly two time intervals in the collapsed state - a behavior that cannot be accounted for by a Markov chain; similarly, discarding information about the state space of a Markov chain usually results in a loss of the Markov property. When it does not, the chain is said to be lumpable. The reader is referred to chapter VI of Kemeny and Snell (1976) for more on this subject. Here, we will not try to find conditions on the life-cycle graph for the strong or weak lumpability of **P**, let us only mention that (1) from Theorem 6.3.2 of Kemeny and Snell (1976), the Markov matrix associated with most models encountered in practice are not strongly lumpable and (2) at the very least, we need the chains associated with P to be weakly lumpable when using π as initial probability distribution, but even this is not easy to turn into simple necessary and/or sufficient conditions on the life-cycle graph.

Therefore, rather than restricting ourself to the (probably very restricted) class of matrix population models that have lumpable ergodic chains, we recognize that, in general, the process obtained by observing the original Markov chain on the set of collapsed classes is non-Markovian, but we seek to approximate it with a Markov chain. It remains to specify what sense we would like to give to this approximation, i.e. which properties we would like the original and the collapsed model to share.

If we want to preserve elasticities, which correspond to the stationary probability distribution on the arcs of **P**, we would like **P**\* and **P** to have compatible stationary probability distribution, i.e. 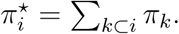. Looking at the calculations that showed that individualistic collapsing preserves the stable stage distribution, we see that this can be achieved by letting

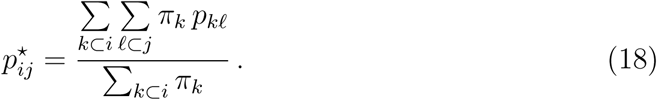
And indeed, it is then immediate that **π**\***P**\* = **π**\*.

Having collapsed (**P**, **w**, λ) into (**P**\*, **w**\*, λ*), all we have to do is reconstruct **A**\* from (**P**\*, **w**\*, λ*) using equation (11). After simplifications, we get

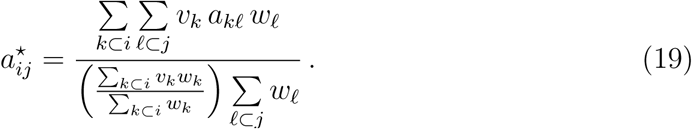
It is easy to check that

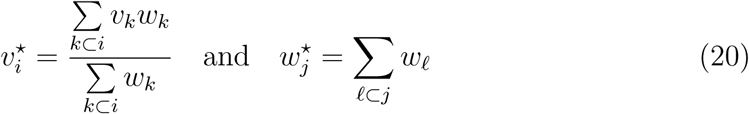
are, respectively, left and right eigenvectors of **A**^*^ associated with λ^*^ = λ. By the same argument as previously, **A**\* being primitive, we conclude that **v**\*, **w**\* and λ* are indeed the vector of reproductive values, the stable stage distribution, and the growth rate of **A**\*, respectively - in accordance with intuition. Similarly, one can check that

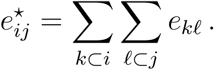
We call this new method for collapsing *genealogical collapsing*. In addition to preserving the growth rate, stable stage distribution, reproductive values and elasticities, this method also has conceptually satisfying properties such as the fact that it commutes with unit conversion, provided that the collapsed conversion factors are defined by

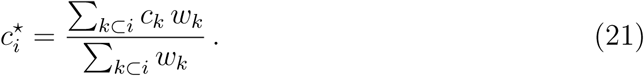
The easiest way to see why this is the case is to remember that, in the (**P**, **w**, λ) decomposition, only w is affected by the choice of units and, clearly, collapsing w commutes with changes of units, since 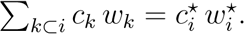.

The fact that genealogical collapsing commutes with unit conversion is conceptually satisfying because there is something arbitrary in the choice of units (even though in the case of population models, the individual usually imposes itself as the natural unit). Moreover, the previous discussion about units allows us to better interpret genealogical collapsing: from formula (19), we can see that it consists in expressing the model in terms of reproductive values, as described by (12), with *c_i_* = *v_i_*, performing individualistic collapsing on the resulting model, and then switching units back to individuals again. This allows us to understand where individualistic collapsing fails: in individualistic collapsing, individuals of different classes are added. But, strictly speaking, an individual of class k is not the same unit as an individual of class *l* ≠ *k*. By contrast, reproductive value is a “central” unit in the sense that only the amount of reproductive value counts, not the individuals carrying it - in accordance with the intuition behind formula (20) for the collapsing of reproductive value. Thus, in genealogical collapsing, adding contributions is made possible by the fact that they have been converted to a common unit first.

Before closing this technical presentation of genealogical collapsing, it should be noted that if the reproductive values of the classes collapsed together are the same 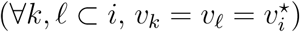, then equation (14) and equation (19) give the same results, that is, genealogical collapsing reduces to individualistic collapsing.

### 4.3 Examples

We close our presentation of genealogical collapsing by working out a few examples that illustrate the difference between individualistic and genealogical collapsing. More specifically, we give three simple models that correspond to very different biological situations but yield almost identical individualistic models.

Consider the projection matrix

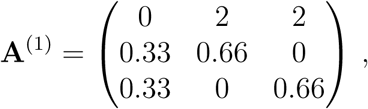
whose life-cycle graph is depicted in figure 2.A. It corresponds to a model with two identical adult classes. The dominant eigen-elements of A^(1)^ are

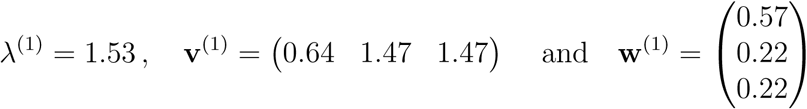
It can be checked from equation (14) that, conforming to intuition,

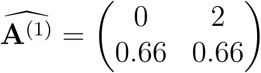
Moreover, 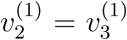, i.e. the classes that are collapsed have the same reproductive value. We have seen that in that case, genealogical collapsing and individualistic collapsing yield the same projection matrix. Therefore, 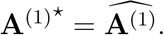.

But now consider the matrix

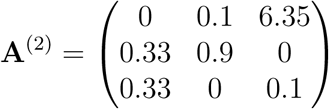
with dominant eigen-elements

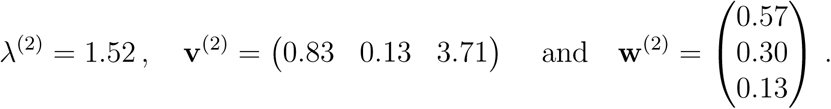
This projection matrix, whose life-cycle graph is depicted in figure 2.B, corresponds to a very different biological situation, where the two adults types are radically different: one has a low fertility and high survival while the other has a high fertility and low survival. This translates into the first one having a lower reproductive value while making-up a bigger proportion of the stable population. Collapsing states 2 and 3 in A^(2)^ gives us:

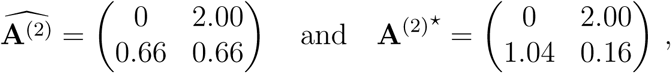
so that different from A^(1)^ though A^(2)^ is, 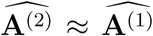. Furthermore, notice that 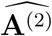 and A^(2)⋆^ are actually quite different, demonstrating that individualistic and genealogical collapsing can give quite different results, even on simple models. More subtly, the bottom left entry of A^(2)*^ also illustrates the fact that, unlike individualistic collapsing, genealogical collapsing does not guarantee that collapsing transitions of weight smaller than 1 yields a transition of weight smaller than 1, forbidding one to interpret the collapsed weight as a survival probability. We will return to this issue in the Discussion.

**Figure 2:**
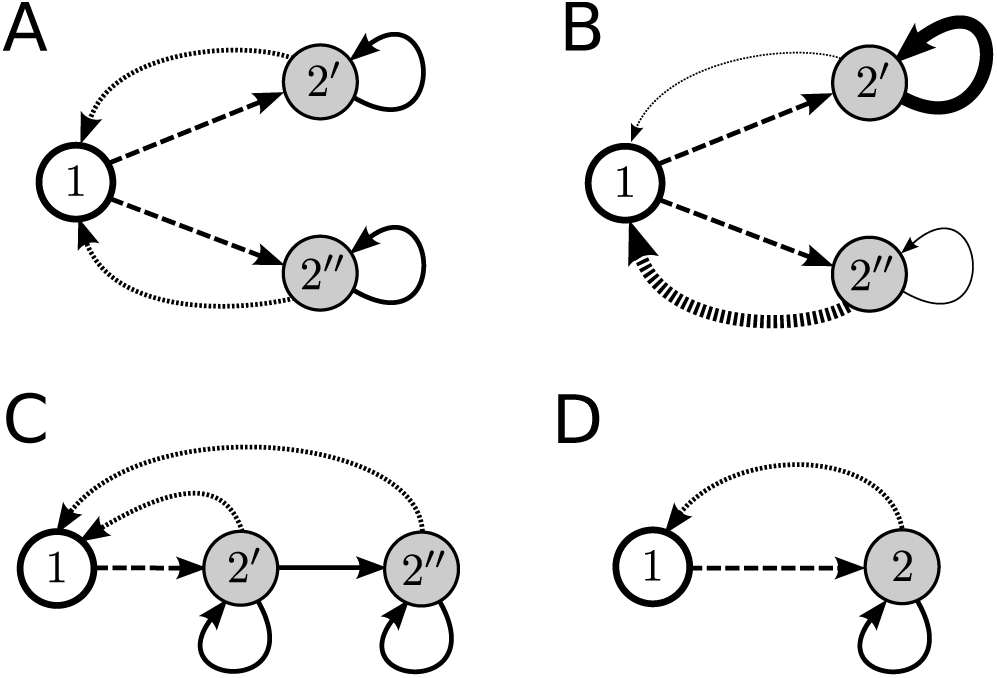
A-C, the life-cycle graphs associated with the projection matrices **A**^(1)^, **A**^(2)^ and **A**^(3)^, respectively. D, the structure of the life-cycle graph obtained by collapsing states 2′ and 2″ in any of these three models.

Finally, consider yet another projection matrix (shown in figure 2.C):

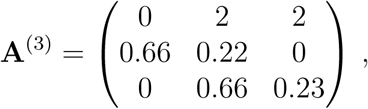
with dominant eigen-elements

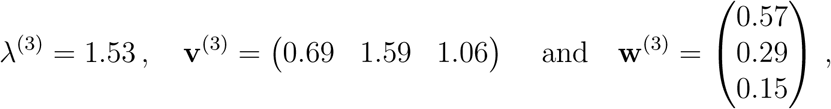
which collapses to

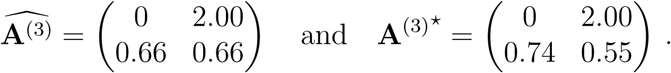
A^(3)^ is yet another example of model that is very different from A^(1)^ and A^(2)^ and yet shares the same individualistically collapsed matrix.

Table 1 gives some classic biological descriptors for each of the models presented above. It shows that the difference between the descriptors of the individualistic model and those of the genealogical one can be non-negligible. Note that, on these particular examples, the net reproductive rate R_0_* of the genealogical model seems to be a better approximation of real value (i.e. the R_0_ of the original model) than the net reproductive rate 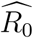 of the individualistic model.

**Table 1:**
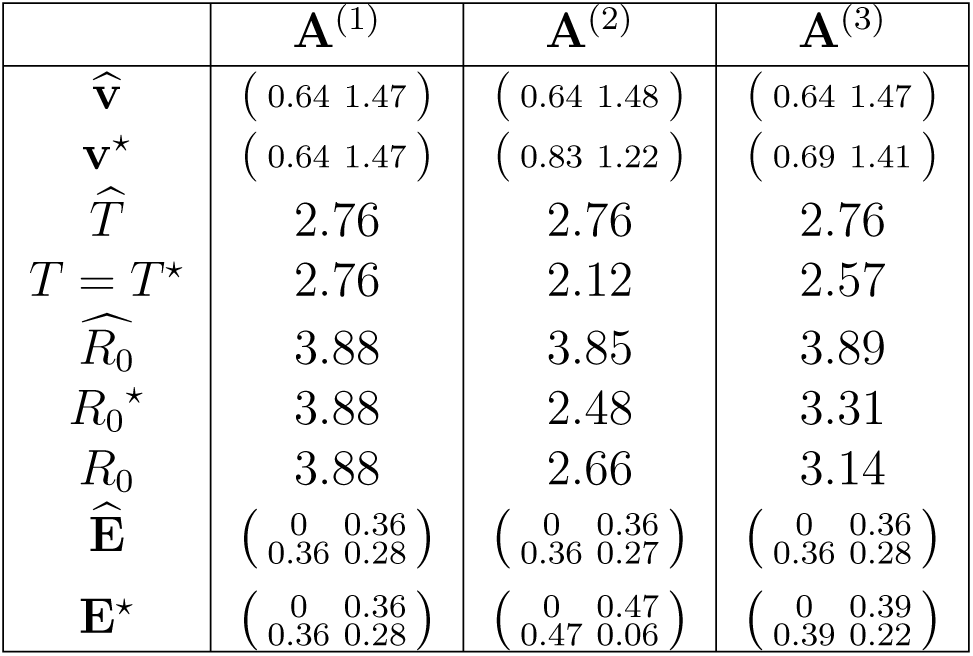
The reproductive value vectors **v**, generation times *T*, net reproductive rates *R_0_* and elasticity matrices **E** of the individualistically collapsed and genealogically collapsed models for each of the three models presented in the main text. Small discrepancies such as the fact that the entries of 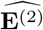 sum to 0.99 are due to the fact that, as in the rest of the section, the results have been rounded to two digits to be displayed.

| | $\mathbf{A}^{(1)}$ | $\mathbf{A}^{(2)}$ | $\mathbf{A}^{(3)}$ |
| --- | --- | --- | --- |
| $\widehat{\mathbf{v}}$ | $\begin{pmatrix} 0.64 & 1.47 \end{pmatrix}$ | $\begin{pmatrix} 0.64 & 1.48 \end{pmatrix}$ | $\begin{pmatrix} 0.64 & 1.47 \end{pmatrix}$ |
| $\mathbf{v}^*$ | $\begin{pmatrix} 0.64 & 1.47 \end{pmatrix}$ | $\begin{pmatrix} 0.83 & 1.22 \end{pmatrix}$ | $\begin{pmatrix} 0.69 & 1.41 \end{pmatrix}$ |
| $\widehat{T}$ | 2.76 | 2.76 | 2.76 |
| $T = T^*$ | 2.76 | 2.12 | 2.57 |
| $\widehat{R}_0$ | 3.88 | 3.85 | 3.89 |
| $R_0^*$ | 3.88 | 2.48 | 3.31 |
| $R_0$ | 3.88 | 2.66 | 3.14 |
| $\widehat{\mathbf{E}}$ | $\begin{pmatrix} 0 & 0.36 \\ 0.36 & 0.28 \end{pmatrix}$ | $\begin{pmatrix} 0 & 0.36 \\ 0.36 & 0.27 \end{pmatrix}$ | $\begin{pmatrix} 0 & 0.36 \\ 0.36 & 0.28 \end{pmatrix}$ |
| $\mathbf{E}^*$ | $\begin{pmatrix} 0 & 0.36 \\ 0.36 & 0.28 \end{pmatrix}$ | $\begin{pmatrix} 0 & 0.47 \\ 0.47 & 0.06 \end{pmatrix}$ | $\begin{pmatrix} 0 & 0.39 \\ 0.39 & 0.22 \end{pmatrix}$ |

## 5. Discussion

Matrix population models are used to calculate the ecological properties of a population (e.g., the population growth rate, the stable stage structure, etc.) from the traits of the individuals composing it (survival probabilities and fertilities). However, the link between the descriptors of the individuals and those of the population can be complex, making it hard to see how some of the ecological descriptors of the population are related. We have suggested an alternative framework to partly solve this problem, in which matrix population models are parametrized by a set of descriptors that capture various aspects of the population dynamics. In this parametrization, rather than directly specify the entries of the population projection matrix (n^2^ degrees of freedom, where n is the number of stages) we instead specify the elasticities, which contain information about the realizations of the life-cycle along the lineages of the genealogy of the population (n^2^ — n degrees of freedom); the stable stage distribution - or, equivalently, the reproductive values - which determines the partitioning of class reproductive value into individuals (n — 1 degrees of freedom); and the asymptotic growth rate, which describes the expansion of the population over time (1 degree of freedom).

The genealogical decomposition follows directly from previously known facts about matrix population models (Demetrius, 1974, 1975; Tuljapurkar, 1982, 1993), but to our knowledge had not previously been recognized as an alternative way of specifying a matrix population model. The main conceptual insight this decomposition offers is that some properties of a structured population depend only on the genealogical properties of the population. Furthermore, these properties are independent of the units we use to keep track of organisms (biomass, individuals, etc.), or equivalently, the form of the stationary stage distribution.

This primary insight is in some ways a generalization of the well-known observation that although demographic events (birth, death, survival) as a rule happen to individuals in various states, to compare the importance of these events for the population dynamics, one needs to “convert” individuals in different states to a common unit by weighing them with their reproductive values (Fisher, 1930). Our results show that one can go one step further, and characterize the dynamics of a population in this common unit separately from the description of how the dynamics manifest themselves in units of individuals.

This kind of separation seems well suited for theoretical studies of matrix population models because, whereas changing one entry of the projection matrix will affect many biological descriptors, changing a parameter in our parametrization will leave many of them unchanged, or affect them in a straightforward way. The parametrization also makes it easy to build projection matrices with prescribed properties. Finally, if clarifies the link between widely used descriptors by showing how some of them can be decoupled.

For instance, there has been a long-going debate about when to use elasticities or sensitivities. Our framework suggests that elasticities should be favored when the properties one is interested in depend only on the genealogy of the population – or, equivalently, when the units used to build the model do not matter; by contrast, when units play a crucial role, using sensitivities seems a better option.

Our framework also helps make sense of recent results on the accumulation of neutral mutations in structured populations (Balloux and Lehmann, 2012; Lehmann, 2014b; Allen et al., 2015; Amster and Sella, 2016). These results show that the neutral substitution rate can be very different than the expected mutation rate of a random individual drawn from the population, in apparent contradiction of classical results in population genetics (Kimura et al., 1968). However, these results appear very natural under our framework because the neutral substitution rate is simply the long-term frequency of mutations along lineages. Thus, the neutral substitution rate depends only on the genealogies and stage-dependent mutation rates and is completely uncoupled from the stationary stage distribution (and hence from the expected mutation rate in a randomly drawn individual).

A third potential use of the genealogical decomposition is to uncover underlying similarities in the life-history of different species that are not apparent in the individual-based projection matrices. Our two-state examples in section 3.2 illustrate this: if the two states are, say, juveniles and adults, populations with very different ratios of juveniles to adults (e.g., high or low x) might nonetheless have similar P matrices. This would mean that the dynamics of lineages are similar in the two populations, and therefore the populations would also have similar elasticities, generation times, and class reproductive values despite their grossly different compositions.

An important limitation of our analysis arises from the deterministic nature of matrix population models. One interpretation of these models is that they describe the dynamics of the expected abundances of the types in a discrete time multi-type branching process (see chapter 5 of Athreya and Ney (1972) for a mathematical presentation of multi-type branching processes, and chapter 16 of Caswell (2001) for more on the link with matrix population models), and all of the ingredients of the genealogical decomposition are available in this framework. For instance, a rich literature exists on the structure and convergence properties of the genealogical Markov chains and their relation to the asymptotic growth rate in the broader context of multi-type branching processes (Hermisson et al., 2002; Georgii and Baake, 2003; Baake and Georgii, 2007; Leibler and Kussell, 2010; Wakamoto et al., 2012; Sughiyama et al., 2015; Kobayashi and Sughiyama, 2015). However, there are also important differences between the determistic and the stochastic setting: whereas in a deterministic setting the choice of units is in some sense arbitrary, in branching processes the choice of some particle as the unit is central to the specification and analysis of the model, as the particles make independent contributions to the next generation.

Another important stochastic generalization of matrix population models arises when considering the joint genealogy of two or more individuals sampled from the population, as in coalescent-based approaches for studying structured populations (e.g., Rousset, 2004). In this context, the choice of units again becomes important because the number of individuals in the class determines the probability of coalescence. Note, however, that the theoretically important unit of individuality in this context is the chromosome rather than the organism. Thus, whereas our analysis depends on the ability to change units while preserving many features of the model, more detailed stochastic models will often have a single natural choice of unit.

We illustrated the practical potential of the genealogical decomposition by showing how it naturally leads to a new solution to the problem of aggregating stages in a matrix population models. This new method, which we dubbed “genealogical collapsing”, has many properties not enjoyed by the standard, “individualistic” method for collapsing, such as the fact that it preserves the reproductive values and the elasticities, and that it is independent of the units of the model.

One obvious application of genealogical collapsing is to allow demographic comparisons between species modeled with different numbers of stages (e.g. Enright et al., 1995). In this case, the applicability of the method is the same as that of individualistic collapsing: any set of classes of any primitive projection matrix can be collapsed thanks to equation (19). Therefore, a natural question arises: when should genealogical collapsing be favored over individualistic collapsing? If the classes collapsed together have the same reproductive value, both methods are equivalent. However, as shown in section 4.3, when the reproductive values of the classes collapsed together differ, the differences can be substantial.

If one is primarily interested in descriptors such as the reproductive values, the elasticities, or descriptors that are expected to be closely related to these, then genealogical collapsing should be used. If by contrast what matters most is the interpretation of the entries of the projection matrix as per capita contributions, then individualistic collapsing seems a better option. In that case, classes should preferentially be grouped based on similarity of the reproductive values, rather than by their similarity in terms of other demographic traits; if the reproductive values of the classes collapsed together differ substantially, then one should be careful about the resulting collapsed model. In any case, it is probably best to compare the results obtained with both methods. Finally, note that if one is only interested in global (that is, non class-specific) quantities such as the net reproductive rate, etc, then these should of course be computed from the original model.

Another possible application of genealogical collapsing is to simplify integral projection models (Easterling et al., 2000; Ellner and Rees, 2006; Merow et al., 2014), where some aspects of the organismal state are continuous, to standard matrix population models containing only discrete states. While current discretization methods often require a very fine grid over the continuous states to provide a good approximation (Zuidema et al., 2010), the theoretical guarantees on the performance of genealogical collapsing suggests that genealogical collapsing might provide more satisfactory results for coarse discretizations. We note, however, that the rigorous analysis of integral projection models is quite technical (Ellner and Rees, 2006) and that the methods described here are only applicable to the extent that a fine discretization provides a sufficiently accurate approximation to the full integral projection dynamics.

A third possible application of collapsing is to assess the impact of within-class heterogeneity in the vital rates. The fact that the vital rates are homogeneous within a class is an underlying assumption of matrix population models, but it is rarely met in practice so it is natural to wonder how this can affects the output of a given model. To estimate this, one can formulate a specific hypothesis about how the heterogeneities are distributed within the classes by thinking of the classes as being composed of several subclasses. The model can then be “extended” into a compatible model, i.e. a model that will give the model from which we started when collapsed using individualistic collapsing. For instance, if the initial model was built by counting individuals and interpreting the entries of the projection matrix as per capita contributions, then the initial model can be seen as the individualistically collapsed model associated with the “true” extended model. Comparing the initial model with the model obtained via genealogical collapsing of the extended model can thus be used to get an idea of how important the discrepancies introduced by the heterogeneities of the vital rates within the classes can be. Note that this scheme has the advantage of allowing for a very selective relaxation of the within-class homogeneity hypothesis, because all the other assumptions on which the study of matrix population models is based are kept.

All of this is in the spirit of the examples presented in section 4.3, where A^(2)^ and A^(3)^ can be viewed as two different hypotheses about the form of heterogeneity in the vital rates of class 2 of model 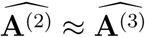. Similarly, most matrix population models encountered in practice are either age-based (Leslie models) or size-based, whereas the vital rates are usually known to depend on both variables. Collapsing can be used to convert between these two types of models, by extending a size-based model into an (age, size)-based model and then collapsing it into an age-only model, as depicted in figure 3. In particular, given a size-based model, once can construct an extended version of this model where the states are (age, size) pairs, but the vital rates are determined entirely by size as in the original model. Collapsing such a model to include only size classes recovers the original model because the vital rates (and hence reproductive values) are determined solely by size. However, reproductive value will typically vary within age classes, and so the results of collapsing to an age only model will depend on the collapsing method used.

**Figure 3:**
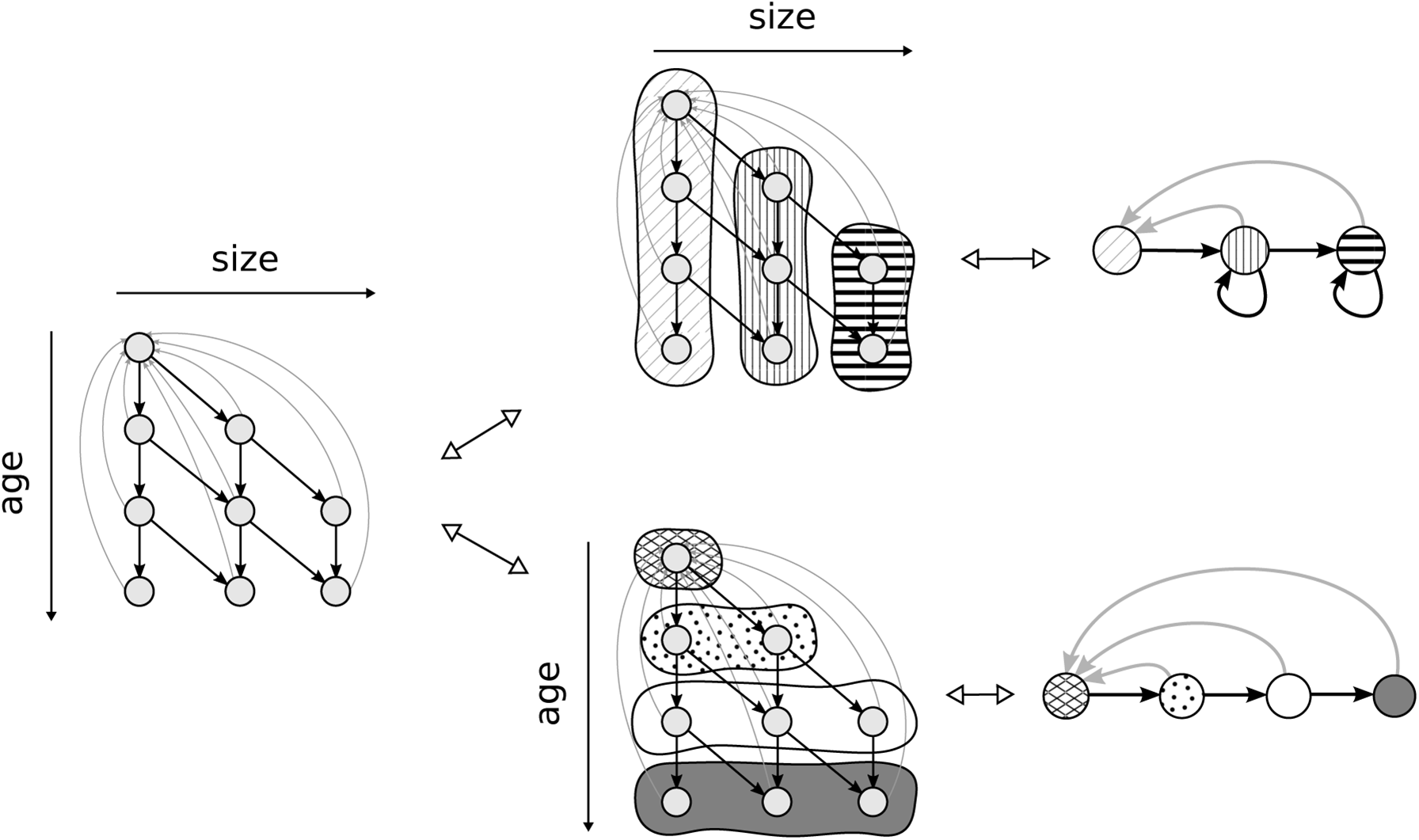
An (age, size)-based model and the corresponding age-based model and size-based model, both obtained by collapsing some of the classes of the (age, size) model. Recovering the original model from the collapsed one requires additional hypotheses about the distribution of vital rates within the classes, made on a per case basis and based on a good knowledge of the organism and of the population being studied. Comparing the individualistic and genealogical model can be used to test how these hypotheses impact the output of the model.

The discussion above highlights an important conceptual implication of our results: since in practice models are built by counting individuals, any matrix population model can be viewed as the individualistically collapsed version of a “real” extended model to which we do not have access. As a result, computing descriptors such as the elasticities or the reproductive values can give incorrect results, even if there is no error in the entries of the projection matrix. In particular, it is a fundamental limitation of matrix population models that they are not resilient to within-class heterogeneities when it comes to computing the reproductive values or the elasticities; by contrast, under stationarity they give consistent estimates of the asymptotic growth rate and of the stable population structure.

A theoretical solution to this problem would be to count contributions in terms of reproductive value rather than in individuals, but this is unlikely to be of much use in practice since reproductive value cannot be measured directly.

Since collapsing is about discarding information from a model, any collapsing method is bound to have shortcomings. We now discuss some of the limitations of genealogical collapsing. The biggest problem of the method is that is does not retain the interpretation of the entries of the population projection matrix as per-capita entries. For instance, we have seen that collapsing transitions that correspond to survival probabilities might yield a transition whose weight is greater than 1, so that it cannot be interpreted as a probability. By contrast, this cannot happen with individualistic collapsing, since the weight obtained by collapsing survival probabilities individualistically is simply the observed average survival probability across the collapsed classes for a population at its stable stage distribution.

However, this defect of genealogical collapsing may not be as much of a drawback as it first appears because great care is needed in the interpretation and analysis of survival probabilities in collapsed models even when using individualistic collapsing. For instance, the classic method most notably developed in Cochran and Ellner (1992) and reviewed in chapter 5 of Caswell (2001) consists in interpreting survival probabilities as the transition probabilities of an absorbing Markov chain that tracks the moves of individuals between the classes of the model during their life-time. However, when viewed in the reduced state-space, the paths taken by individuals under the original model will typically not be Markovian, violating a key assumption of the Cochran and Ellner (1992) method. While individualistic collapsing does optimally preserve the one-step survival probabilities, other descriptors calculated by the Cochran and Ellner (1992) method will generally not be preserved. On the other hand, genealogical collapsing results in transition rates that cannot be interpreted as survival probabilities but which nonetheless maintain several key global properties of the population dynamics such as the generation time and reproductive values, and which optimally preserve the one-step transition probabilities of the stationary genealogical Markov chains defined by the matrices P and Q. Furthermore, while entries of the genealogically collapsed projection matrix cannot be interpreted as per-capita contributions, their interpretation is actually straightforward from a different point of view: since it does not make sense to add-up individuals from different classes, we express all contributions in terms of reproductive values, sum them, and then convert back the result in terms of individuals by using the collapsed reproductive value as the conversion factor.

It should also be noted that even though individualistic collapsing is the *de facto* standard for collapsing projection matrices, in some specific situations other methods have been suggested. For instance, Yearsley and Fletcher (2002) proposed a method that preserves the asymptotic growth rate, the stable stage distribution and the generation time when collapsing adjacent stages in what are often called size-based models (cf figure 3). Genealogical collapsing also enjoys all of these properties, but in addition preserves the reproductive values and the elasticities, and is more general in that it can be used to collapse any group of classes in any primitive matrix population model.

Although genealogical collapsing is interesting in its own right, it is important to see that that its properties flow fundamentally from its close relationship with the reparameterization of matrix population models described here. Because this parametrization allows the different global properties of the model to be altered independently from each other, each can be collapsed separately and thereby optimally preserved. By clarifying the relationship between various global features of matrix population models, we hope that this reparameterization will spur further innovation in the inference and analyses of these models.

## Acknowledgements

We thank Laurent Lehmann and two anonymous reviewers for helpful comments.

